# DupLoss-2: Improved Phylogenomic Species Tree Inference under Gene Duplication and Loss

**DOI:** 10.1101/2024.09.05.611565

**Authors:** Rachel Parsons, Mukul S. Bansal

## Abstract

Accurate species tree reconstruction in the presence of widespread gene duplication and loss is a challenging problem in eukaryote phylogenomics. Many phylogenomics methods have been developed over the years to address this challenge; these range from older methods based on gene tree parsimony to newer quartet-based methods. In this work, we introduce improved software for gene tree parsimony-based species tree reconstruction under gene duplication and loss. The new software, DupLoss-2, uses an improved procedure for computing gene losses and is far more accurate and much easier to use than its previous version released over a decade ago. We thoroughly evaluate DupLoss-2 and eight other existing methods, including ASTRAL-Pro, ASTRAL-Pro 2, DISCO-ASTRAL, DISCO-ASTRID, FastMulRFS, and SpeciesRax, using existing benchmarking data and find that DupLoss-2 outperforms all other methods on most of the datasets. It delivers an average of almost 30% reduction in reconstruction error compared to iGTP-Duploss, the previous version of this software, and a 10% reduction compared to the best performing existing method. DupLoss-2 is written in C++ and is freely available open-source.

## 1 Introduction

Phylogenomics, which involves the use of genome-scale sequence data for phylogenetic inference, has emerged as one of the most powerful techniques for accurate species tree reconstruction. However, complex evolutionary processes such as gene duplication, gene loss, horizontal gene transfer, incomplete lineage sorting, and hybridisation can create complex patterns of incongruence between the evolutionary histories of different genomic loci, confounding phylogenomic analyses (Maddison 1997, Steenwyk et al. 2023). In particular, gene duplication and loss (*duplication-loss* for short) is one of the most important drivers of gene family and genome evolution in eukaryotes (Ohno 1970, Zhang 2003), but many phylogenomics approaches are unable to directly use gene families affected by duplication-loss (Kapli et al. 2020, Smith and Hahn 2021). To address this limitation, many methods have recently been developed for duplication-loss phylogenomics. Such methods can use multicopy gene families as well as gene families not universally present in the species under consideration, allowing them to make more efficient use of available genomic data (Smith and Hahn 2021).

Nearly all methods for duplication-loss phylogenomics take as input a collection of gene trees, corresponding to hundreds or thousands of gene families, and estimate a species tree from them based on some objective function. Several existing methods are based on gene tree-species tree reconciliation frameworks that explicitly model duplication-loss events. Such methods include DupTree (Wehe et al. 2008), iGTP-DupLoss (Chaudhary et al. 2010), DynaDup (Bayzid et al. 2013), PHYLDOG (Boussau et al. 2013), and SpeciesRax (Morel et al. 2022). Other methods do not explicitly model duplication-loss and instead rely on evolutionary-process-agnostic notions of distance between gene trees and species trees. Examples include NJ(st) (Liu and Yu 2011) and DISCO-ASTRID (Willson et al. 2021), based on estimating distances between pairs of taxa, and MulRF (Chaudhary et al. 2014b) and FastMulRFS (Molloy and Warnow 2020), based on generalised Robinson-Foulds distance between multi-labeled trees. Recently, methods based on quartet analysis have also emerged as a promising approach for duplication-loss phylogenomics. Such methods include ASTRAL-Pro (Zhang et al. 2020), DISCO-ASTRAL (Willson et al. 2021), and ASTRAL-Pro 2 (Zhang and Mirarab 2022).

In this work, we make two key contributions. First, we provide substantially improved software for duplication-loss phylogenomics. The new software, named *DupLoss-2*, is based on the duplication-loss reconciliation model (Goodman et al. 1979, Page 1994, Mirkin et al. 1995, Eulenstein and Vingron 1998, Zhang 1997, Slowinski and Page 1999, Ma et al. 2000) and uses a new procedure for computing gene losses that results in substantially improved species tree accuracy compared to iGTP-DupLoss (Chaudhary et al. 2010), the previous version of this software. It also provides a streamlined command-line user interface that is far easier to use for large-scale phylogenomic analysis than iGTP-DupLoss. And second, we use the extensive benchmarking dataset of Chaudhary et al. (2014a) to characterize the performance of DupLoss-2 and eight other state-of-the-art methods, including ASTRAL-Pro (Zhang et al. 2020), ASTRAL-Pro 2 (Zhang and Mirarab 2022), DISCO-ASTRAL (Willson et al. 2021), DISCO-ASTRID (Willson et al. 2021), MulRF (Chaudhary et al. 2014b), FastMulRFS (Molloy and Warnow 2020), and SpeciesRax (Morel et al. 2022), across a wide range of evolutionary and experimental conditions.

Our experimental analysis identifies DupLoss-2 as the most accurate method overall among all evaluated methods, yielding an approximately 10% reduction in reconstruction error on average compared to the second-best method. Importantly, DupLoss-2 significantly outperforms iGTP-Duploss, improving species tree accuracy by an average of almost 30%, across all tested evolutionary and experimental conditions including different rates of gene duplication and loss, number of taxa, number of input gene trees, gene tree error rates, and gene tree incompleteness rates. DupLoss-2 also outperforms all other methods on 19 of the 27 individual datasets, performing especially well on datasets with medium or high rates of duplication-loss as well as on datasets with smaller numbers of gene trees. Our results also provide valuable and sometimes surprising insights into the performance of other methods; for example, we find that ASTRAL-Pro can fail to run on datasets with fewer gene trees or larger numbers of taxa, that the accuracy of FastMulRFS can degrade sharply as the number of taxa increases, and that ASTRAL-Pro, ASTRAL-Pro 2, DISCO-ASTRAL, and FastMulRFS are almost always more accurate than DISCO-ASTRID, MulRF, iGTP-DupLoss, and SpeciesRax. Overall, our results indicate that DupLoss-2 can lead to substantial improvement in species tree reconstruction accuracy on datasets where gene duplication and loss are the primary drivers of gene family evolution.

## 2 Description

### 2.1 Overview of DupLoss-2

DupLoss-2 is based on the gene tree parsimony approach (Guigó et al. 1996, Page and Charleston 1997, Page 1998) and builds upon the older duplication-loss phylogenomics software packages DupTree (Wehe et al. 2008) and iGTP-DupLoss (Chaudhary et al. 2010). At the most basic level, DupLoss-2 takes as input a collection of gene trees and seeks a species tree that minimizes the total duplication-loss reconciliation cost, i.e., invokes the fewest total number of gene duplications and losses to reconcile all the input gene trees. Since the corresponding computational problem is likely NP-hard (see Bansal and Eulenstein (2013) for a nuanced discussion on what is known about the complexities of different variants of this problem), DupLoss-2 uses heuristics to infer the species tree. Specifically, DupLoss-2 first constructs an initial candidate species tree using a greedy step-wise leaf-addition heuristic, and then executes a subtree pruning and regrafting (SPR) based local search heuristic to iteratively improve the current species tree. The local search heuristic terminates when a better species tree cannot be found in the SPR neighborhood of the current species tree, and DupLoss-2 outputs the resulting species tree. Such SPR-based local search heuristics are widely used for phylogenetic tree search, such as in RAxML (Stamatakis 2014), DupTree (Wehe et al. 2008), FastME (Lefort et al. 2015) and other methods. For improved scalability, DupLoss-2 specifically implements the fast SPR search algorithms of Bansal and Eulenstein (2013) that can find the species tree with lowest duplication-loss reconciliation cost in an SPR search neighborhood within Θ(*mn*) time (compared to Θ(*mn*^2^) time required for a brute force search), where *n* represent the numbers of leaves in the species tree and *m* the total number of leaves in all the input gene trees

The input gene trees can be either rooted or unrooted and can, optionally, be assigned different weights to control their relative contributions to the total reconciliation cost (see below). If any of the input gene trees are unrooted, DupLoss-2 automatically reoptimizes their root positions periodically during the local search, similarly to iGTP-DupLoss (Chaudhary et al. 2010).

*Additional features*. DupLoss-2 also includes several useful features to facilitate rigorous phylogenomic analyses. These features include (i) differential gene tree weighting, allowing for input gene trees to be assigned different user-specified weights based on their quality, size, etc., (ii) the ability to impose structural constraints on the inferred species tree, allowing for evaluation of different evolutionary hypotheses and for improved species tree accuracy, (iii) implicit randomness in the tree search, such that multiple executions of DupLoss-2 on the same input can be used for improved thoroughness and accuracy and for sampling multiple optimal or near-optimal species trees; a Python script is provided for easy execution of multiple replicate runs on the same input dataset, (iv) options for post-hoc analysis of input gene trees, allowing for quick insight into the reconciliation of individual input gene trees with the inferred species tree, (v) the ability to use a user-provided initial species tree for the local search in lieu of generating an initial species tree internally, allowing for greater control over the search space, and (vi) the ability to use user-provided random number generator seeds, allowing for exact replication of any analysis.

### 2.2 Key improvements

DupLoss-2 improves upon its predecessor in two important ways. First, DupLoss-2 uses a different calculation for the number of losses that is better attuned to the characteristics of modern phylogenomics datasets. And second, DupLoss-2 provides a streamlined and easy-to-use command-line interface that allows for easy parallelisation and is more in line with how phylogenomics inference programs are typically used today. We elaborate on these points below.

Improved calculation of losses. The previous version of DupLoss was released as part of the iGTP software package over a decade ago (Chaudhary et al. 2010). We refer to this previous (and original) version of DupLoss as *iGTP-DupLoss*. At the time, only a small number of species had fully sequenced genomes and, as a result, the gene families used for phylogenomic analyses were frequently incomplete. In other words, even though a species may be included in the phylogenomic analysis, not all of its genes may have been sequenced/sampled, leading to considerable uncertainly about whether a gene was truly absent from that species’ genome or whether the absence was an artifact of incomplete genome sequencing. To account for this uncertainty, the loss calculation implemented in iGTP-DupLoss first trims the species tree being evaluated to only the taxa represented in the gene tree currently under consideration. Essentially, the loss calculation of iGTP-DupLoss errs on the side of undercounting losses rather than risk overcounting them. However, when whole-genome datasets are used for phylogenomic analyses, as is now the norm, the number of losses can be estimated more accurately by not trimming the species tree for each gene tree. Thus, in most cases, it is no longer appropriate to trim the species tree for computing losses. Consequently, the new loss calculation implemented in DupLoss-2 does not trim the species tree when computing losses.

Figure 1 illustrates the difference between computing losses on trimmed versus untrimmed species trees. As our experimental results show, this relatively small change in how losses are computed significantly impacts the tree search and leads to drastic improvements in species tree inference accuracy (see experimental results in next section). Importantly, and somewhat surprisingly, we find that DupLoss-2 significantly outperforms iGTP-DupLoss even on datasets with incomplete gene sampling (see next section).

**Figure 1.**
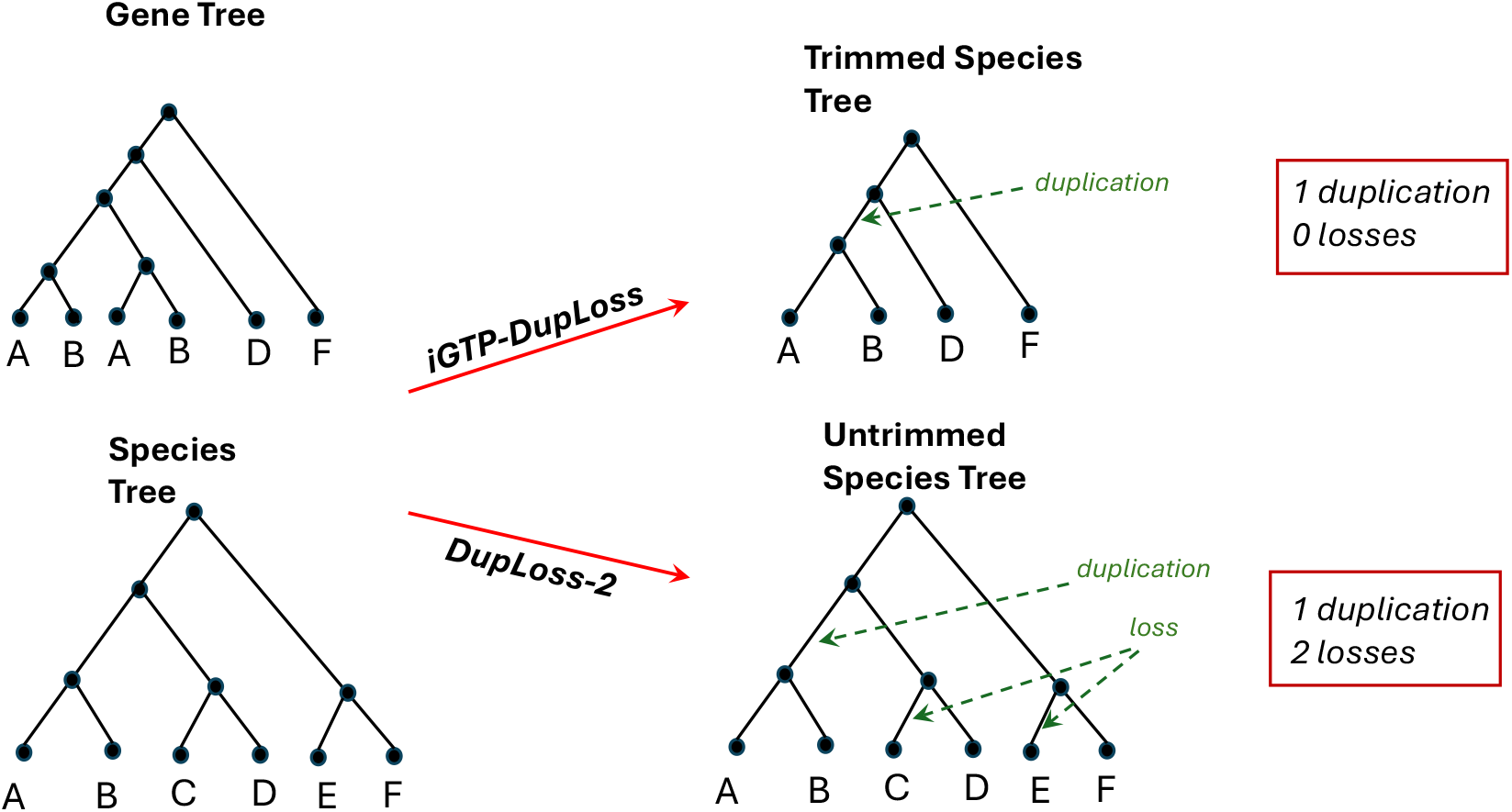
Calculating losses on trimmed versus untrimmed species trees. The figure shows the number of duplications and losses inferred by iGTP-DupLoss (top right) and DupLoss-2 (bottom right) for the gene tree and species tree shown in the left column. The loss calculation implemented in iGTP-Duploss first reduces (or trims) the species tree to only the species represented in the gene tree being reconciled. For this example, iGTP-DupLoss therefore invoke one duplication and no losses, as shown. The new loss calculation implemented in DupLoss-2 uses the full, untrimmed species tree and therefore infers 1 duplication and 2 losses (along the species tree edges shown by the dotted green arrows).

#### Improved user interface

DupLoss-2 provides a streamlined command line interface that makes it far easier to use and much more amenable to rigorous, large-scale phylogenomic analyses. In contrast, iGTP-DupLoss only provides a graphical user interface, which makes its use impractical in many situations. For instance, iGTP-DupLoss makes it difficult to use compute servers or high performance computing resources to run long jobs or perform more thorough phylogenomic analyses, and it cannot be used within automated computational analysis pipelines. DupLoss-2 is designed to be easy to use, with only the name of the input file containing gene trees as a required parameter, and has easily parseable output to ease integration with computational pipelines. Such command-line interfaces have been found to be easier to use in most cases, with popular phylogenomics software like DupTree (Wehe et al. 2008) and ASTRAL (Mirarab et al. 2014) employing similar interfaces.

## 3 Experimental Evaluation

To demonstrate the utility of DupLoss-2, we perform a thorough evaluation of the accuracies of DupLoss-2 and many other state-of-the-art methods using an extensive simulated benchmarking dataset. This benchmarking dataset was previously used in a study comparing the accuracies of several older methods, including DupTree (Wehe et al. 2008), iGTP-DupLoss (Chaudhary et al. 2010), MulRF (Chaudhary et al. 2014b), NJ(st) (Liu and Yu 2011), and PHYLDOG (Boussau et al. 2013). That previous study identified iGTP-DupLoss and MulRF as the two best methods for duplication-loss phylogenomics (Chaudhary et al. 2014a). Since then many newer methods have been developed for duplication-loss phylogenomics, with ASTRAL-Pro (Zhang et al. 2020), ASTRAL-Pro 2 (Zhang and Mirarab 2022), DISCO-ASTRAL (Willson et al. 2021), DISCO-ASTRID (Willson et al. 2021), FastMulRFS (Molloy and Warnow 2020), and SpeciesRax (Morel et al. 2022) being among the most promising. Our experimental analysis includes these six newer methods, as well as the older methods iGTP-DupLoss and MulRF. Thus, a total of nine methods, including DupLoss-2, were evaluated.

### 3.1 Description of dataset

The benchmarking dataset of Chaudhary et al. (2014a) was carefully simulated to evaluate the effect of five different aspects of phylogenomic studies on the accuracy of duplication-loss phylogenomics methods. These are (i) number of taxa (species) in the analysis, (ii) rates of duplication and loss (D/L), (iii) gene tree inference error, (iv) number of input gene trees, and (v) level of incompleteness of input gene trees. Specifically, the benchmarking dataset consists of multiple individual datasets, each representing a different combination of simulation parameters, and with each individual dataset consisting of 20 replicates. Specific settings for these five simulation parameters appear below, with default values underlined:

1. **Number of taxa:** 10, 50, <u>100</u>, 500.
2. **D/L rates:** low, medium, and high. These rates correspond to gene duplication and loss rates of 0.002, 0.004, and 0.008 events/gene per myr, respectively, estimated from primate data.
3. **Gene tree inference error:** error-free and <u>error-prone</u>. The error-prone gene trees were inferred using RAxML and simulated sequences of length 500nt.
4. **Number of input gene trees:** <u>1000</u>.
5. **Gene tree incompleteness:** <u>0</u>, 0-25, 25-50, and 50-75 percent of sequences deleted from gene trees. Further details on how these datasets were originally simulated appear in Chaudhary et al. 2014a. In order to more fully assess the impact of gene tree error and number of input gene trees, we created additional datasets with higher rates of gene tree inference error and datasets consisting of only 500 and 250 gene trees. The gene trees with higher error rates were inferred by trimming down the 500nt sequence alignments to 300nt and then applying RAxML, and the datasets with 500 and 250 gene trees were created by simply selecting 500 or 250 gene trees at random from the 1000-gene tree datasets. Thus, the corresponding simulation parameter settings can be updated as follows:
6. **Gene tree inference error:** error-free, <u>error-prone</u>, and high-error-prone. The error-prone and high-error-prone gene trees were inferred using RAxML and simulated sequences of length 500nt and 300nt, respectively.
7. **Number of input gene trees:** <u>1000</u>, 500, 250.

### 3.2 Overall experimental setup

While DupLoss-2 and iGTP-DupLoss can take as input both rooted and unrooted gene trees, the other methods included in this study all use unrooted gene trees as input. Thus, all analyses were performed using only unrooted input gene trees. Species tree reconstruction accuracy was calculated by comparing inferred unrooted species trees to corresponding ground truth species trees using the widely used (unrooted) normalized Robinson-Foulds distance (NRFD) (Robinson and Foulds 1981). NRFD takes values between 0 and 1 and represents the fraction of non-trivial splits in the inferred species tree that are not present in the ground truth species tree. Thus, lower NRFD values indicate better reconstruction accuracy.

Each method was executed using its default parameter settings and was run exactly once on each input. Reported results are averaged across all 20 replicates in any individual dataset.

### 3.3 Results

#### Overall performance trends

We first evaluated the impact of the two most important evolutionary parameters, D/L rate and number of taxa (or species tree size), on the accuracies of the nine different methods. These datasets used default values for the other parameters (i.e., 1000 error-prone gene trees as input). Figure 2 shows the results of this evaluation for all twelve combinations of D/L rate and number of taxa. These results reveal many important insights into the relative performance of different methods for duplication-loss phylogenomics. First, we find that DupLoss-2 substantially outperforms iGTP-DupLoss on each of the twelve datasets, with DupLoss-2 showing an approximately 28% reduction in NRFD compared to iGTP-DupLoss when averaged across all D/L rates and numbers of taxa. Second, DupLoss-2 outperforms all other methods on eight of the twelve datasets, with either ASTRAL-Pro, DISCO-ASTRAL, or FastMulRFS outperforming DupLoss-2 on the other four datasets. In general, DupLoss-2 tends to outperform all other methods on the datasets with medium and high D/L rates, while one of the other methods tends to outperform DupLoss-2 on the low D/L rate datasets. Third, somewhat surprisingly, ASTRAL-Pro generally slightly outperforms its successor, ASTRAL-Pro 2, on these datasets. However, we also found that ASTRAL-Pro failed to run on over half of the 500-taxon datasets, while ASTRAL-Pro 2 was able to run on all datasets. And fourth, we find that DISCO-ASTRID, MulRF, iGTP-DupLoss, and SpeciesRax generally show worse accuracy than the other methods on these datasets. Overall, when averaged across all twelve datasets (all D/L rates and numbers of taxa), DupLoss-2 shows the lowest average NRFD at 0.244, with DISCO-ASTRAL, ASTRAL-Pro 2, and FastMulRFS emerging as the second-, third-, and fourth-best methods with NRFDs of 0.286, 0.287, and 0.293, respectively. DupLoss-2 remains the best overall method even after excluding the results for 500-taxon datasets (on which ASTRAL-Pro and MulRF could not be successfully executed), with an average NRFD of 0.204 and with ASTRAL-Pro showing the second-best average NRFD at 0.241.

**Figure 2.**
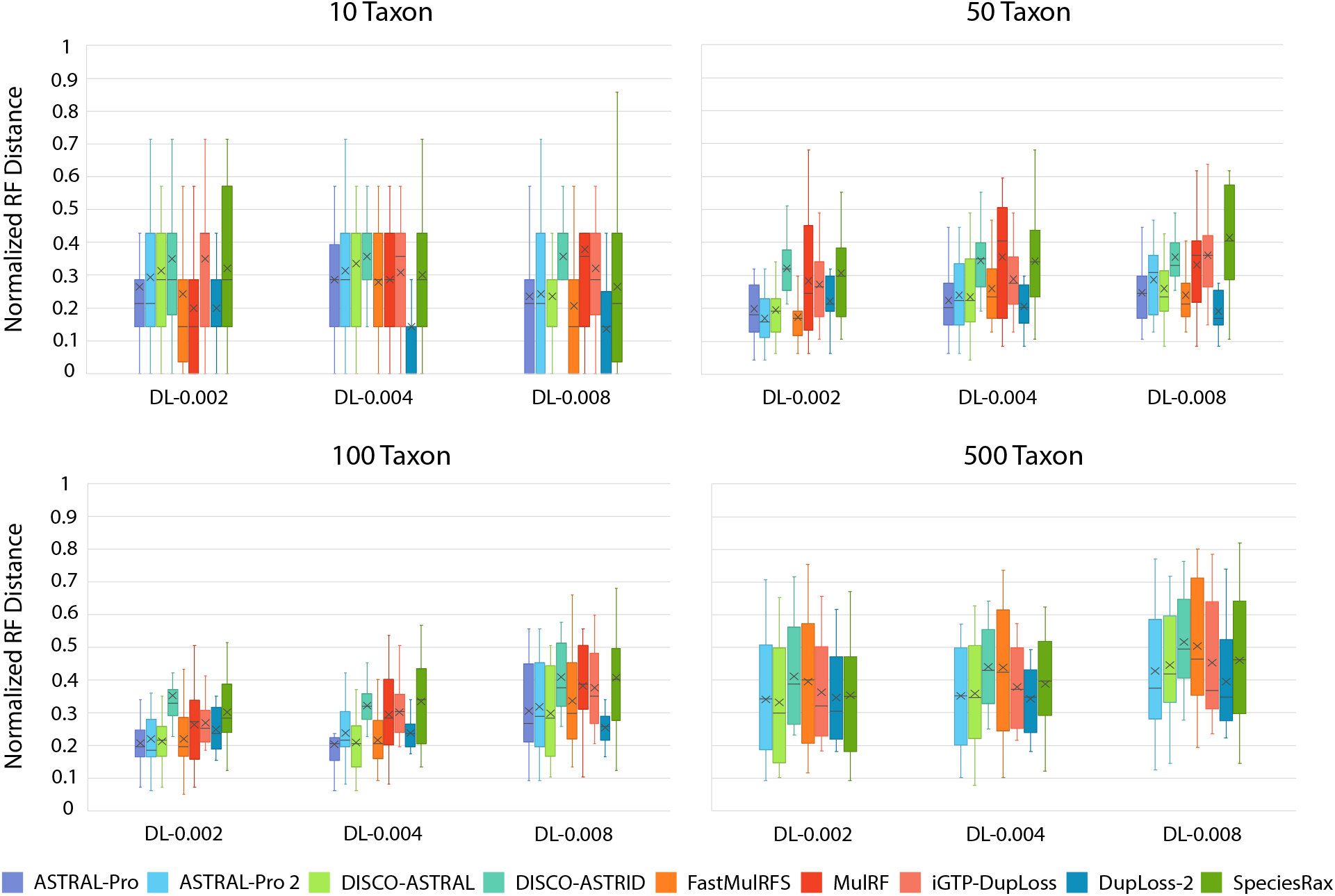
Overall accuracy trends for different methods. Species tree reconstruction accuracies are shown for the nine different methods on datasets with different D/L rates (low, medium, and high D/L rates; columns within each boxplot) and different numbers of taxa (10, 50, 100, and 500 taxa; different boxplots). Average NRFDs are marked by “x” within each box. Results are shown for datasets consisting of 1000 error-prone gene trees. Lower NRFD values imply greater reconstruction accuracy. ASTRAL-Pro and MulRF are excluded from the results for 500-taxon datasets; ASTRAL-Pro failed to run on 33 of the 60 500-taxon datasets, while MulRF was excluded because of lengthy execution times (multiple days per 500-taxon dataset).

#### Impact of gene tree error

To assess the impact of gene tree error on the different methods, we evaluated the accuracies of the different methods on datasets with error-free (true) gene trees and datasets with higher gene tree error rates (high-error-prone) than the default error-prone datasets. Figure 3 shows these results. On the datasets with error-free gene trees, we find that DupLoss-2 drastically outperforms all other methods across all D/L rates, inferring nearly error-free species trees with an average NRFD of 0.014 across the three D/L rates. ASTRAL-Pro, the second-best method on these datasets, shows a much higher average NRFD of 0.032. While error-free gene trees represent an unrealistic scenario in practice, these results shed light on how well each method can be expected to work under ideal conditions. On the datasets with higher error rates (high-error-prone), we find that FastMulRFS shows the best performance on low D/L datasets, DISCO-ASTRAL shows the best performance on medium D/L datasets, and DupLoss-2 shows the best performance on high D/L datasets. Almost all methods show a significant reduction in accuracy on these datasets compared to the datasets with default error (error-prone; see plot for 100 taxa in Figure 2). We note that ASTRAL-Pro failed to run on 2 of the 20 high D/L datasets and its high D/L results were averaged over only the 18 successful runs. Overall, DupLoss-2 shows the lowest average NRFD of 0.322 across all D/L rates, with FastMulRFS showing the second-best performance with an average NRFD of 0.332. These results suggest that DupLoss-2 can substantially outperform all other methods on datasets with high D/L rates, even with gene trees of poor quality.

**Figure 3.**
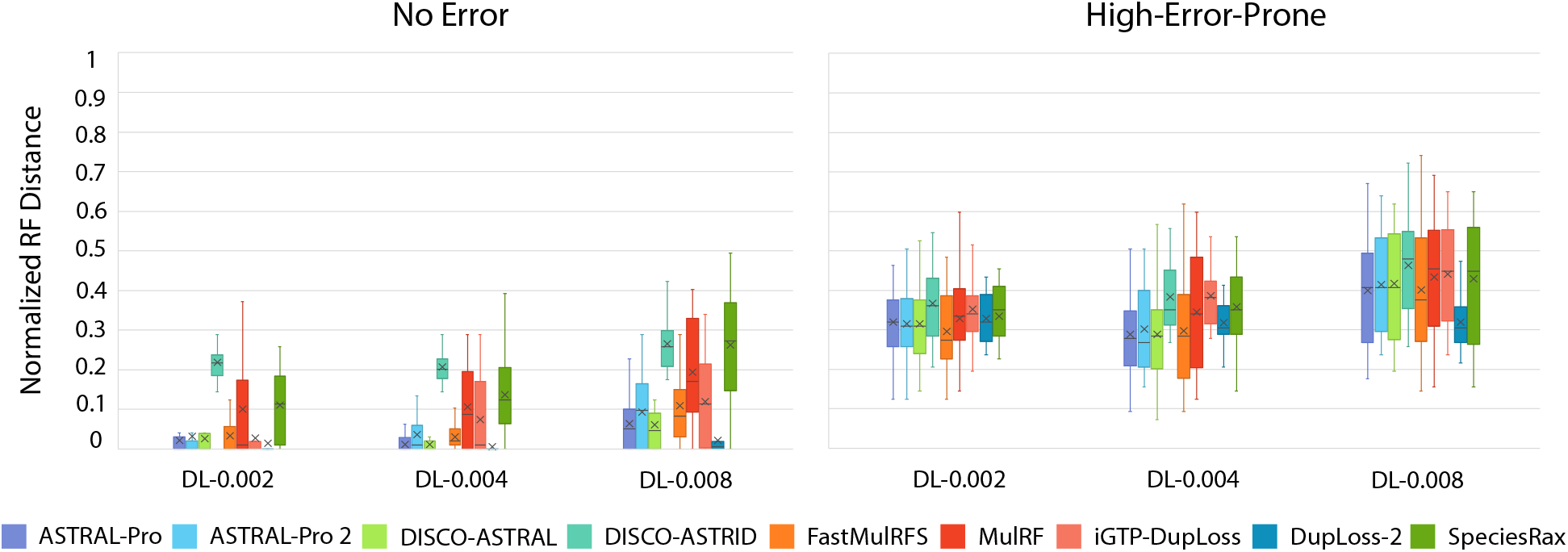
Impact of gene tree error. Species tree reconstruction accuracies are shown for the nine different methods on datasets with error-free gene trees (left boxplot) and datasets with higher error rates than the default error-prone datasets (right boxplot). Average NRFDs are marked by “x” within each box. All datasets were generated using default values for the other simulation parameters, i.e., 100 taxa and 1000 input gene trees. Lower NRFD values imply greater reconstruction accuracy. ASTRAL-Pro failed to run on 2 (out of 20) of the high D/L rate datasets for high-error-prone gene trees and its accuracy was averaged over the 18 successful runs.

#### Impact of number of gene trees

We next assessed the impact of number of gene trees on species tree reconstruction accuracy. Figure 4 shows the results of this assessment. We find that DupLoss-2 continues to be the best overall method when the number of gene trees is reduced, outperforming all other methods for medium and high D/L rates on both the 500- and 250-gene tree datasets. For low D/L rate, FastMulRFS performs best on the 500-gene tree datasets and DupLoss-2 performs best on the 250-gene tree datasets. These results show that DupLoss-2 can make efficient use of the available information and lead to substantially improved accuracy compared to other methods on datasets with smaller numbers of gene trees. For instance, the average NRFD of DupLoss-2 across all D/L rates on the 250-gene tree datasets is 0.387, while the average NRFD for the second-best method, ASTRAL-Pro, is 0.425.

**Figure 4.**
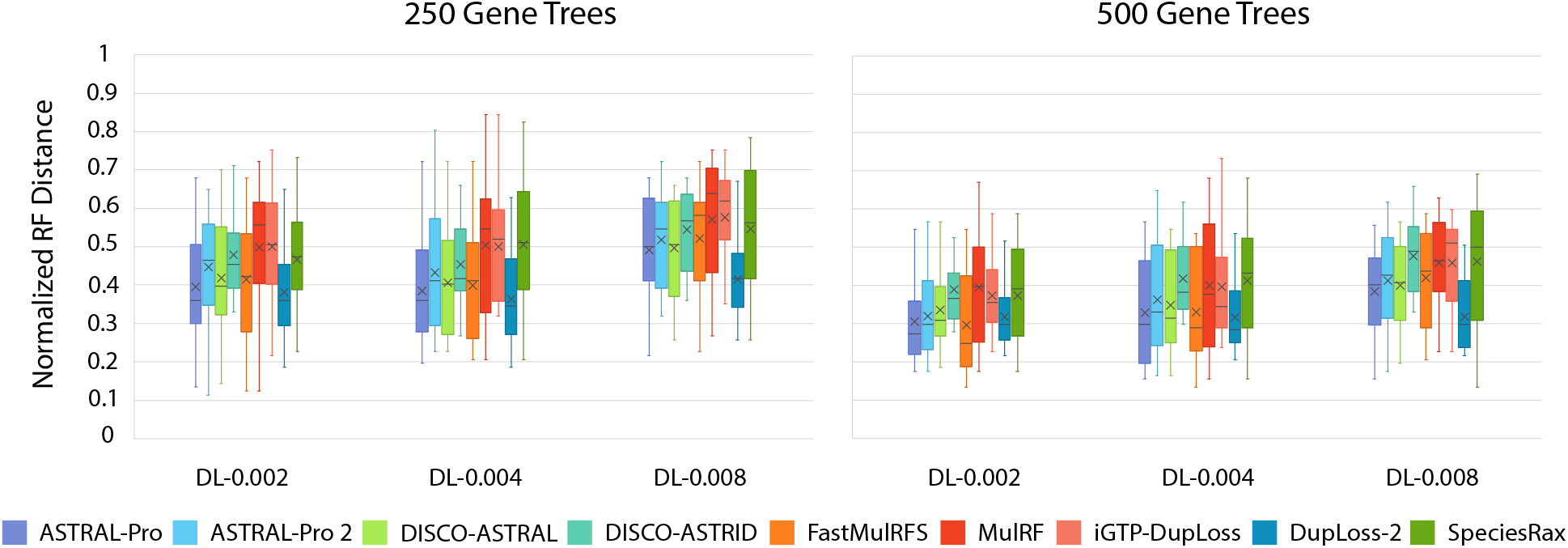
Impact of reduced number of gene trees. Species tree reconstruction accuracies are shown for the nine different methods on datasets with 500 gene trees (left boxplot) and 250 gene trees (right boxplot). Average NRFDs are marked by “x” within each box. All datasets were generated using default values for the other simulation parameters, i.e., 100 taxa and error-prone gene trees. Lower NRFD values imply greater reconstruction accuracy. ASTRAL-Pro failed to run on 2 (out of 20) of the high D/L rate, 500-gene tree datasets and on 1 (out of 20) of the low D/L rate, 250-gene tree datasets, and its accuracy was averaged over its successful runs.

We note that ASTRAL-Pro failed to run on 2 (out of 20) of the high D/L rate, 500-gene tree datasets and on 1 (out of 20) of the low D/L rate, 250-gene tree datasets and its results were averaged over the remaining successful runs.

#### Impact of gene tree incompleteness

Finally, we assessed the impact of incomplete sampling of genes, or gene tree incompleteness, on the ability of the different methods to infer species trees. As Figure 5 shows, DupLoss-2 continues to be the best performing method overall. In particular, DupLoss-2 outperforms all other methods for the datasets with 25-50% sequences removed and 50-75% sequences removed and is the second-best method for the datasets with 0-25% of sequences removed. Interestingly, DupLoss-2 continues to substantially outperform iGTP-DupLoss across all three rates of sequence removal. This result is surprising since the loss calculation of iGTP-DupLoss is specifically designed to deal with incomplete gene trees. Overall, these results show that DupLoss-2 is highly robust to incomplete gene sampling.

**Figure 5.**
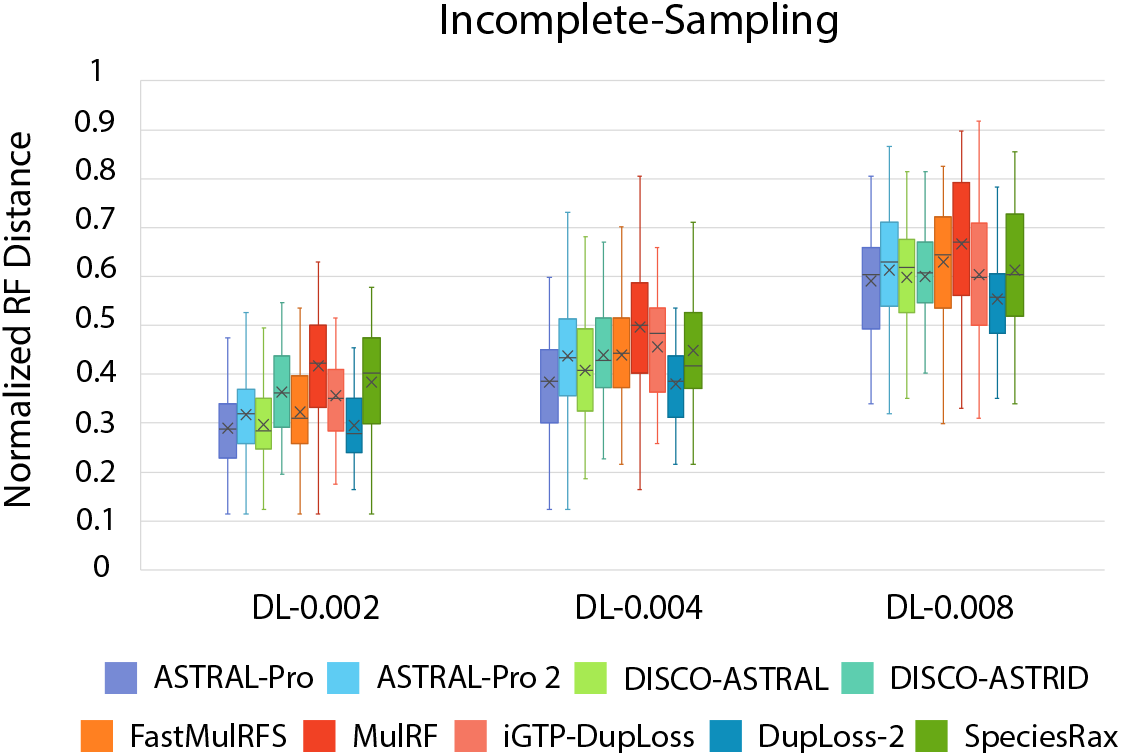
Impact of gene tree incompleteness. Species tree reconstruction accuracies are shown for the nine different methods on datasets with three different levels of gene tree incompleteness. These datasets correspond to the following simulation parameter settings: 100-taxa, low D/L rate, and 200 error-prone gene trees (Chaudhary et al. 2014a). Average NRFDs are marked by “x” within each box, and lower NRFD values imply greater reconstruction accuracy.

#### Running time analysis

We also measured average running times of the different methods on datasets with increasing numbers of taxa (Table 1). All tested methods were scalable to datasets up to 500 taxa, the largest dataset size we evaluated, but there was wide variability in running times. DISCO-ASTRAL is the fastest method, requiring only about 8 seconds on average on the 500-taxon dataset, and MulRF is the slowest, requiring about 64 hours on the same 500-taxon datasets. All methods were run using a single core of an Intel Xeon 2.1 GHz processor. Overall, ASTRAL-Pro, ASTRAL-Pro 2, DISCO-ASTRAL, DISCO-ASTRID, and FastMulRFS are the fastest methods, requiring no more than a few minutes on average on the 500-taxon datasets, while SpeciesRax, DupLoss-2, iGTP-DupLoss, and MulRF are considerably slower, requiring approximately 7, 36, 46, and 64 hours, respectively. These results show that DupLoss-2, though much slower than the fastest methods, is still highly scalable and can be applied to datasets with hundreds of taxa within a few hours.

**Table 1.** Ruuning times in hours, minutes, and seconds are shown for the nine different methods on medium D/L rate datasets with 1000 error-prone gene trees and different numbers of taxa. Running times are not shown for ASTRAL-Pro since it failed to run on a large fraction of the 500-taxon datasets. All methods were executed using a single core of a 2.1 Ghz Intel Xeon processor with 64 GB of RAM.

| Method | 10 taxa | 50 taxa | 100 taxa | 500 taxa |
| --- | --- | --- | --- | --- |
| ASTRAL-Pro | 0h:0m:3s | 0h:0m:13s | 0h:0m:24s | - |
| ASTRAL-Pro 2 | 0h:0m:0.20s | 0h:0m:5s | 0h:0m:19s | 0h:5m:21s |
| DISCO-ASTRAL | 0h:0m:1s | 0h:0m:10s | 0h:0m:23s | 0h:4m:43s |
| DISCO-ASTRID | 0h:0m:0.50s | 0h:0m:1s | 0h:0m:1s | 0h:0m:8s |
| FastMulRFS | 0h:0m:0.69s | 0h:0m:19s | 0h:1m:9s | 0h:14m:36s |
| MulRF | 0h:0m:11s | 0h:11m:50s | 1h:5m:18s | 63h:42m:55s |
| iGTP-DupLoss | 0h:0m:2s | 0h:3m:5s | 0h:27m:2s | 46h:5m:10s |
| DupLoss-2 | 0h:0m:2s | 0h:3m:40s | 0h:35m:19s | 36h:25m:59s |
| SpeciesRax | 0h:0m:8s | 0h:1m:24s | 0h:4m:25s | 6h:47m:43s |

## 4 Discussion

In this work, we introduce the DupLoss-2 software for duplication-loss phylogenomics and perform thorough evaluation of DupLoss-2 and eight other methods using a large benchmarking dataset. DupLoss-2 uses an improved loss calculation and provides a user-friendly, streamlined command line interface that is amenable to parallelisation and is easy to include in bioinformatics analysis pipelines. In our experiments, DupLoss-2 shows the highest overall species tree reconstruction accuracy (lowest average NRFD) among all evaluated methods, being especially effective for datasets with medium or high rates of duplication and loss as well as for datasets with lower numbers of gene trees. Notably, DupLoss-2 significantly outperforms the previous version of this software, iGTP-Duploss, under all tested evolutionary and experimental conditions. The predecessors of DupLoss-2, including iGTP-DupLoss and DupTree, are widely used for phylogenomic analyses of eukaryotic species (see, e.g., Ceron-Romero et al. 2022, Marletaz et al. 2023, Gerdol et al. 2020, Alioto et al. 2020, Green et al. 2014, Dohm et al. 2014, Vonk et al. 2013, Garcia-Mas et al. 2012, Katz et al. 2012 for some recent and other high profile applications of these methods), and our results show that DupLoss-2 can lead to substantially improved species tree accuracy over these existing software.

Our results also demonstrate how several newer duplication-loss phylogenomics methods, such as ASTRAL-Pro, ASTRAL-Pro 2, and DISCO-ASTRAL, provide a clear improvement in accuracy over older methods such as MulRF and iGTP-DupLoss. On the other hand, our results also suggest that some newer methods, such as SpeciesRax and DISCO-ASTRID, may not be well-suited for duplication-loss phylogenomics, at least when using their default parameter settings. FastMulRFS also performed well, but our results suggest that its accuracy degrades sharply as the number of taxa in the analysis increases. Overall, our experimental results show that using DupLoss-2 can lead to significant improvements in species tree reconstruction accuracy on datasets where gene duplication and loss are the primary drivers of gene family evolution and gene tree discordance.

Our results suggest that while evolutionary-process-agnostic methods such as ASTRAL-Pro, DISCO-ASTRAL, ASTRAL-Pro 2, and FastMulRFS perform well, they are likely to be outperformed by methods like DupLoss-2 that explicitly model evolutionary processes. Thus, it may help to develop more sophisticated methods that explicitly model many of the underlying evolutionary processes. For example, in many eukaryotic datasets, especially those where closely related species are analyzed, gene-tree/species-tree discordance may result from a combination of gene duplication and loss, incomplete lineage sorting (ILS), hybridisation, and other evolutionary phenomena. Several complex reconciliation models that jointly handle duplication-loss and ILS already exist (Rasmussen and Kellis 2012, Wu et al. 2013, Paszek et al. 2021, Mishra et al. 2024), but most suffer from high computational complexity and have not yet been utilized for phylogenomics. We note that *joint* modeling of multiple processes seems necessary for improved accuracy, since phylogenomics approaches that consider multiple simple reconciliation costs, but do not model the underlying processes jointly, often fail to outperform approaches based directly on simple reconciliation models (De Oliveira Martins et al. 2014, Saha et al. 2024).

## Availability

DupLoss-2 source code, precompiled executables for Linux and Mac, user manual, etc., are freely available through GitHub at the following URL: https://github.com/Bansal-CompBioLab/DupLoss-2. The benchmarking dataset of Chaudhary et al. (2014a) used in this work is freely available from Dryad at the following URL: http://dx.doi.org/10.5061/dryad.mr3g6.

## Acknowledgements

This work was supported in part by NSF award IIS 1553421 to MSB.

## Notes

https://github.com/Bansal-CompBioLab/DupLoss-2

### Competing Interest Statement

The authors have declared no competing interest.

